# Modeling and Tracking of Heterogeneous Cell Populations via Open Multi-Agent Systems

**DOI:** 10.1101/2025.09.02.673711

**Authors:** A. Tramaloni, A. Testa, S. Avnet, S. Massari, G. Di Pompo, N. Baldini, G. Notarstefano

## Abstract

Understanding cellular dynamics represents a critical challenge in biomedical research. Optical microscopy remains a key technique for observing live-cell behaviors *in vitro*. This paper introduces an enhanced cell-tracking algorithm designed to address dynamic changes in cell populations, including mitosis, migration, and cell-cell interactions, even within complex co-culture models. The proposed method involves three main steps: 1)modeling the movements and interactions of different cell types in co-culture experiments via tailored open multi-agent systems; 2)identifying parameters via real data for a multi-agent, multi-culture framework; 3) embedding the model within an Extended Kalman Filter, to predict the dynamics of heterogeneous cell populations across video frames. To validate the approach, we used a novel dataset involving the interplay between tumor and normal cells, namely osteosarcoma and mesenchymal stromal cells, respectively. This dataset offers a challenging and clinically relevant framework to track cell proliferation and study how cancer cells evolve and interact with stromal cells within their surroundings. Performance metrics demonstrated the effectiveness of the algorithm over state-of-the-art methodologies, highlighting its ability to track heterogeneous cell types, capture their interactions, and generate the estimated cell lineage tree.

## Introduction

In recent years, the study of cells has gained interest in the control community. Recent works include [1], [2]. The main idea behind these works is to leverage system theoretical approaches to model and control cell populations. These contributions span macro areas like cybergenetics [3] and multicellular systems modeling and simulation [4]. Relevant to our work is the application of multi-agent system paradigms to capture the emergent behaviors observed in heterogeneous cell populations, where individual cells can be modeled as interacting agents. Experimental data for these tasks are typically obtained from optical microscopy videos. Thus, to understand cell behaviors and their interactions, a preliminary task is to extract cell trajectories over time automatically. Biological cell tracking microscopy videos are thus becoming a fundamental task within broad application domains. Among the others, the tracking of tumor interaction with normal stromal cells *in vitro* is crucial to study tumor aggressiveness and drug resistance [5]. Among different techniques, optical microscopy is widely used to retrieve cellular images across different time instants [6].

### A. Related Works

We structure the literature review into three main parts. First, we examine existing cell tracking algorithms. Second, we explore multi-agent approaches in systems and synthetic biology, including how biological principles inspire the design of multi-agent robotic systems. Finally, we review system and control theory methods for analyzing and regulating cell population dynamics.

Multi-Cell Tracking (MCT) is a well-established problem in the literature, with applications ranging from cybergenetics to drug discovery [7], [8]. Given the significant interest in MCT, various datasets and challenges have been developed, such as the Cell Tracking Challenge [9], which serves as a benchmark for MCT algorithms on *in vitro* monoculture experiments. This study focuses on a novel co-culture dataset representing a new challenging scenario compared to other publicly available datasets.

Recent MCT algorithms [10], [11] combine convolutional neural networks (CNN) with recurrent units for segmentation and tracking in microscopy images. Deep learning-based methods include the use of CNNs with Viterbi algorithms [12], Faster R-CNN for multi-feature fusion [13], graph neural networks for temporal modeling [14], and single CNNs for joint segmentation and tracking [15]. Alternative approaches employ Markov random field models with spatial particle filters [16] or variational optical flow techniques [17]. Unlike these black-box methods, our approach combines predictive filtering with tailored multi-agent models that exploit cell-to-cell interactions to improve motion prediction. Moreover, to the best of the authors’ knowledge, existing methods focus primarily on monoculture experiments, whereas co-culture settings accounting for cell population heterogeneity remain relatively uncommon.

Next, works connecting multi-agent system and systems biology are reviewed. Multi-agent modeling has been explored as a framework for designing emergent collective behaviors in synthetic biology [18], such as studying synchronized bacterial populations [19], and coordinating multi-robot systems inspired by chemotaxis principles [20]. For open multi-agent systems, the theoretical framework in [21] provides a basis for studying systems with time-varying state spaces. In our work, we build on similar concepts to model the dynamics of the considered “open” cell system, which is subject to population changes over time.

As for approaches that analyze cell behaviors and model biological interactions, several works often leverage concepts from systems theory, identification, and control. For example, [22] models tumor cell behavior using ordinary, partial, or stochastic differential equations, but does not address single-cell dynamics. Control-oriented approaches are prevalent in related literature, including feedback regulation of cellular populations [23], [24], control of populations endowed with synthetic toggle switches [1], [2], eco-evolutionary models of cancer under therapy [25], and adaptive control of nonlinear cancer-immune dynamics [26]. Overall, these works primarily emphasize control strategies. In contrast, our work focuses on model design, identification, and prediction of cell behaviors.

### B. Contributions

The contributions of this paper are as follows. We propose a novel algorithm, called Heterogeneous Open Multi-Cell EKF-based Tracking (HEOM-EKF), for tracking cells in time-lapse *in vitro* multi-culture experiments, with a specific focus on co-culture settings. HEOM-EKF addresses the following key limitations of existing MCT approaches.

#### Modeling heterogeneous cell populations

Most tracking algorithms are designed for monoculture settings and do not explicitly model interactions among different cell types. *Our contribution:* we model heterogeneous cell populations as an open multi-agent system [27] and propose a class of parametric interaction models that explicitly capture both inter- and intra-type dynamics (e.g., chemotaxis-like behaviors [28]) among cells such as MSC and osteosarcoma. The framework naturally handles switching dynamics in the population over time (e.g., due to mitosis). We focus on these two cell types because, based on several studies—both our own and those of others over the past decades—their interaction has been shown to play a crucial role in inducing aggressiveness and drug resistance in tumor cells [29]–[32].

#### Interpretability of motion prediction models

Many state-of-the-art methods rely on deep learning predictors, which often act as black boxes and provide limited insight into the underlying biological mechanisms. *Our contribution:* we design a structured open multi-agent dynamical model whose parameters are identified from real cell trajectories via tailored system identification procedures.

#### Handling of population change events

Many MCT methods do not robustly distinguish mitosis events from cells entering or exiting the field of view. *Our contribution:* we formulate an Extended Kalman Filter tailored to open multi-agent systems, equipped with switching label sets and dimensionality adaptation. This framework explicitly accounts for mitosis, entry, and exit events, enabling the reconstruction of the cell lineage over time.

In addition, we construct a new annotated dataset from raw time-lapse *in vitro* co-culture experiments involving normal mesenchymal stromal cells and transformed osteosarcoma cells. Unlike most publicly available benchmarks, which focus on monocultures, this dataset provides a challenging and clinically relevant testbed for tracking heterogeneous interacting cell populations and for validating the proposed modeling and identification framework.

A conference version of this work appeared in [33], where we proposed a monoculture cell-tracking algorithm based on a standard multi-agent formulation and evaluated it on a public dataset. That preliminary study assumed a constant number of cells and did not account for mitosis or cells entering and exiting the camera field of view. In contrast, this journal version addresses a more challenging co-culture scenario and leverages a private, unique dataset. Methodologically, we introduce an open multi-agent framework that handles multiple agent (cell) types and explicitly models dynamic population changes. Furthermore, the proposed algorithm reconstructs the cell lineage tree, enabling a deeper analysis of cell interactions and temporal evolution.

### C. Paper Organization

The paper is organized as follows. In Section II, we introduce the cell-tracking problem and outline the essential components of multi-object tracking algorithms. Section III formalizes the open multi-agent system, generalized for multi-culture scenarios, and presents the model identification strategy. In Section IV, we describe the proposed HEOM-EKF algorithm. Section V describes the dataset and its use for training and testing, analyzes the identified multi-cell model parameters, and presents numerical experiments that assess the performance of the proposed algorithm.

## II. Multi-Cell Tracking Problem Formulation

In this work, we consider a cell-tracking problem in a biological scenario in which the goal is to track multiple cells of different types in time-lapse sequences. Throughout the paper, we consider video frames in which the number of cells changes over time. These variations may result from biological processes such as mitosis, and/or from cells entering or exiting the microscope camera field of view (FOV). In this section, we formalize the fundamental concepts behind cell-tracking algorithms. We will delve into more details in Section III, in which we model the cell population as an open multi-agent system and propose suitable formulations for their dynamics. These dynamical models will be a key component of the proposed cell-tracking strategy in Section IV.

The proposed framework follows the widely adopted Track-by-Detection paradigm, in which object detections are first obtained independently on each frame and then linked over time by a dedicated tracking module. In this setting, the main building blocks are: *i*) object detection, *ii*) motion prediction, and *iii*) data association. A schematic of this procedure is depicted in Figure 1. Object detection refers to the identification and localization of cells within images or video frames. Modern object detection methods often employ trained neural networks specifically for this task [34]. Motion prediction aims to estimate future positions of detected cells by combining their past trajectories with current detection measurements. This step is essential for ensuring track continuity and handling gaps or occlusions in the detections. Common techniques include Kalman filters, particle filters, and, most commonly, deep learning-based predictors. However, neural network predictors act as black boxes, thus limiting interpretability, and require large datasets, which are rare in this field. Instead, our approach models multi-cell behaviors explicitly and employs a closed-form solution for the identification of parameters, ensuring reliability and data efficiency. Data association involves linking detected cells across consecutive frames to preserve their identities over time. This process requires matching detections from different frames and assigning them to the correct cell track. Motion models and similarity metrics are frequently used to assess the likelihood of accurate associations.

**Figure 1.**
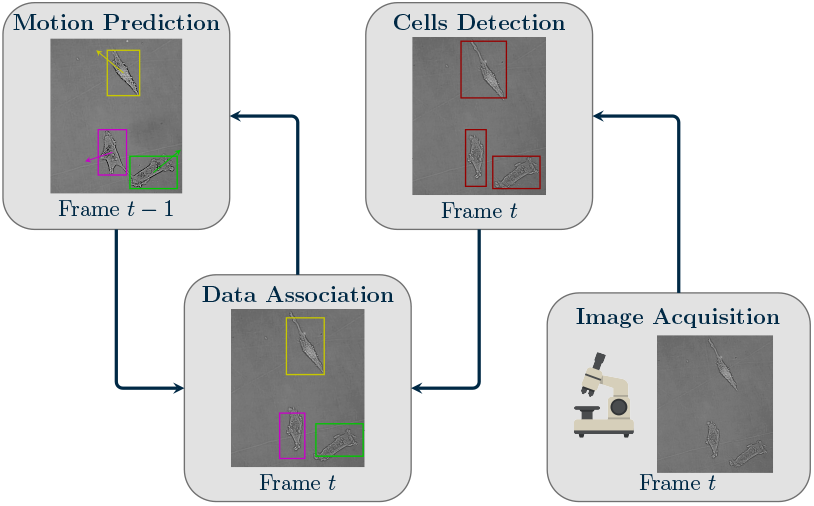
Scheme of a generic multi-cell tracking algorithm

## III. Heterogeneous Cell Populations AS Open Multi-Agent Systems

In this section, we first formalize cell dynamics as an open multi-agent system. Then, we introduce a tailored class of parametric multi-agent terms to describe the interactions between heterogeneous cell populations within a general multi-culture scenario. These dynamical terms constitute the multi-agent model that will be then adopted in the proposed cell-tracking algorithm. The model design draws inspiration from a well-established line of research on bio-inspired multi-agent systems [28]. In particular, we extend to an open multi-agent framework neighboring terms motivated by swarm-like approaches, as in [35] and [36], that describe both positional and morphological cell state behaviors. The latter terms are weighted by a set of parameters to be identified through data collected from time-lapse experiments.

### A. Open Multi-Cell System Formalization

We model mitosis as an event where a parent cell splits into two new daughter cells. In this process, the parent cell is removed from the system and two new cells appear in its place. The latter cells retain the same type (i.e. agent type) as their parent. Moreover, to maintain distinct identities of cells over time, each cell is assigned a unique identifier label *i*. We consider a discrete-time framework, where *t* ∈ *𝒯* ⊆ *∕* denotes the time index over an arbitrary horizon 𝒯 (infinite or finite). In the context of a multi-culture experiment, we restrict attention to a finite time horizon, namely *𝒯* = {1, …, *T*} . Here, *t* will be the discrete-time variable representing the frame at which the system is observed, and with *T*∈ ℕ \ {∞} denoting the final time instant at which the experiment concludes. At any time *t* ∈ *𝒯*, we denote by *N*^*t*^ ∈ *∕* the number of cells present within the microscope FOV. Additionally, we define the subset of time instants 𝒯_*s*_⊂𝒯 corresponding to switching times at which events related to cell population changes occur, i.e. when the number of cells within the observed frame changes, indicating a switch in the system dynamics. We denote the number of cells entering and exiting the heterogeneous cell population within the microscope FOV at time *t* by 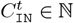 and 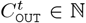, respectively. Let *L*^*t*^ ⊂ *∕* be the set of labels identifying all the cells present in the scene at time *t*. Consequently, we introduce its subsets 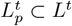 and 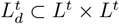 containing the labels of parent and daughter cells, respectively. Based on that, mitoses occurrences can be described through time-dependent sets of label pairs, identifying parent cells splitting into daughter cells at the current time. Formally, mitoses occurring at frame *t* are represented by

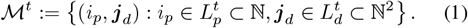

Instead, new cell entries into the camera field, at time *t*, are captured by the set 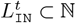, such that

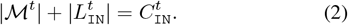

Similarly, cell exiting from the camera FOV at time *t* are described by the label set 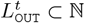, verifying

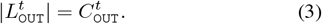

Note that 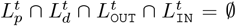, but for different time instants these sets may have non-null intersections. Moreover, at least one of these sets is non-empty at times *t*∈ 𝒯_*s*_. The total number of cells at any time *t*, previously introduced as *N*^*t*^, evolves according to

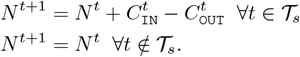

Moreover, the dynamics of the label set *L*^*t*^ can then be expressed as

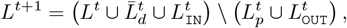

where 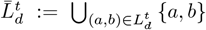 is the set of the individual elements from daughter cells label pairs.

Building upon the concepts introduced above, the following subsection presents the proposed open multi-agent parametric model.

### B. Open Multi-Agent Parametric Model

We now introduce the open multi-cell system that models the cell dynamics for the tracking purpose. In this framework, each generic agent *i* represents a cell. The state of agent *i* at time *t* and its neighbor set are denoted by 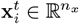 and 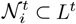, respectively. To support a general multi-culture scenario in which different cell populations are interacting, we consider Γ ∈ 𝒩distinct agent types, each representing a different cell type. Thus, each cell type will be associated a parameter 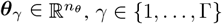,. We denote

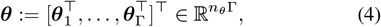

the stack of the parameter vectors for each cell type. The constant parameters ***θ*** will be learned from experimental data.

Let 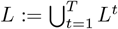 denote the set of all agent labels across time. Then, each agent *i* ∈ *L* is associated with a parameter vector 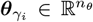, determined by its type *γ*_*i*_ ∈ {1, …, Γ}. Based on that, agent dynamics are modeled according to the discrete-time formulation

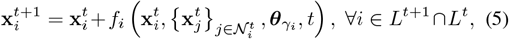

where the function *f*_*i*_ captures the dynamics of agent *i* based on its current state, the states of its neighboring agents, and its type-based parameters 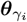 . This formulation accounts for both cell state and type interactions, allowing the model to represent heterogeneous behaviors within a multi-culture experimental setting.

We propose coordinates of a Minimum Enclosing Rectangle (MER) as agent state within the proposed MCT algorithm. This representation of cell morphology is defined as

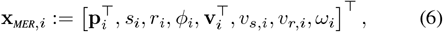

where **p**_*i*_, **v**_*i*_∈ ℝ2 denote the center position and velocity of the cell, *s*_*i*_, *r*_*i*_ ∈ ℝ its area and aspect ratio, and *v*_*s,i*_, *v*_*r,i*_∈ ℝ their respective rates of change. The orientation angle *ϕ*_*i*_ ∈ [0, 2*π*] specifies the rotation of the MER relative to a fixed reference axis, and *ω*_*i*_ ∈ ℝ is the angular velocity. This state representation is especially suitable for tracking cells with fusiform or anisotropic morphologies, as it captures orientation and aspect more faithfully than the standard bounding box format. Moreover, it leads to more informative outputs at the end of the tracking pipeline, supporting better discrimination between different types based on shape dynamics. However, standard performance metrics for tracking algorithms are typically designed for axis-aligned bounding boxes. As a result, evaluating tracking accuracy on the state format in (6) requires custom metrics that are not always readily available or comparable with prior work. To facilitate benchmarking and compatibility with existing evaluation frameworks, our algorithm also supports a simplified state representation based on axis-aligned bounding boxes. Formally,

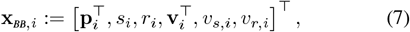

where the state variables retain the same semantics as in (6), but omit orientation and angular velocity. This compact version is especially useful for performance evaluation and comparison with other methods, while still capturing essential information about cell shape and motion.

Based on these two representations, the neighbors of cell *i* at time *t* in (5) can be defined as

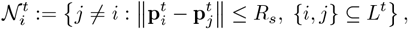

where *R*_*s*_ is a sensing radius that defines the region of space in which other cells are perceived. Here, we consider *R*_*s*_ as a priori fixed quantity (not learnable).

Building on the generic formulation of the open multi-agent model in (5), we now specify the individual terms governing the dynamics of each state component. For better readability, we first define subgroups of the whole state

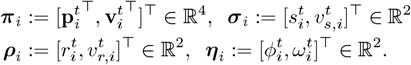

Then, for the MER state format, the dynamics of cell *i* of type *γ*_*i*_ are given by

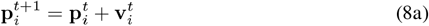

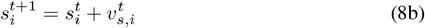

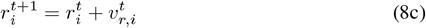

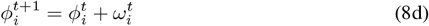

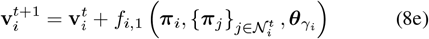

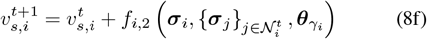

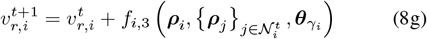

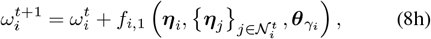

where

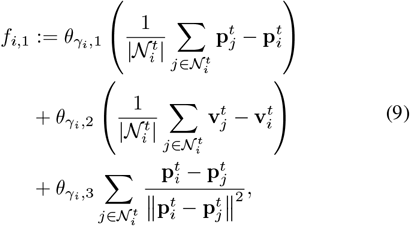

captures the tendency to cohesion, alignment, and repulsion behaviors, while the shape-related terms

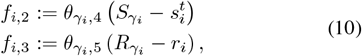

model the tendency to maintain a certain target area 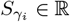 and target aspect ratio 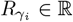, considering cells of type *γ*_*i*_. Here, 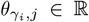 with *j* = 1, …, 5 represent the learnable parameters associated with cells of type *γ*_*i*_.

These multi-cell dynamics are inspired by engineering multi-agent models. Specifically, the terms in (9) follow swarm-based approaches and incorporate chemotaxis-inspired principles [35], [36]. In contrast, the morphological terms in (10), which govern area and aspect ratio dynamics are novel. These terms are designed based on our dataset analysis, where *S*_*γ*_ and *R*_*γ*_ are derived from morphological statistics (see Section V). Together, these components capture a biologically motivated tendency for each cell type to maintain its characteristic shape over time. Finally, we note that for the standard bounding box state defined in (7), the dynamics can be fully described using only equations (8a)-(8c) and (8e)-(8g).

### C. Identification of Model Parameters

We now present the proposed approach for identifying parameters within the multi-agent dynamics discussed in Section III-B. Here, we consider a generic formulation with Γ agent populations captured by parameters 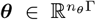, thus providing a flexible tool scalable to different experimental frameworks. Therefore, we propose an identification method tailored to open multi-cell systems, like those described in Section III-A.

The goal is to identify the parameters and obtain estimates that fit a training set of real cell trajectories extracted from *in vitro* co-culture experiments. Consider a dataset consisting of *T* image frames with the relative cell segmentations and tracks. From the latter, for each frame *t* we have access to 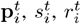 and 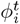 for all cells *i*. From these, we compute the relative velocities 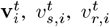 and 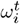 using differentiation methods. We tested various differentiation methods commonly used in system identification from time series data, such as finite differences, smoothed finite differences, Savitzky-Golay filtering, spline interpolation, and Kalman-based filtering, which are also discussed in the context of data-driven model identification (e.g., in [37]). In the experimental tests, we adopted a Kalman-based differentiation approach, which provided smoother estimates and led to smaller residuals in the subsequent model identification phase. This approach allows us to approximate the necessary time derivatives despite the intrinsic stochasticity and noise in biological cell dynamics and aligns with established practices in the literature.

Since we defined the dynamics in equation (5) as a weighted sum of multi-agent state terms, and imposed sensing radius *R*_*s*_ to be not learnable, we end up having a system with linear dependence in ***θ***. Therefore, by properly redefining the system we can construct and solve a typical least squares (LS) problem. The solution to the LS problem is obtained by solving

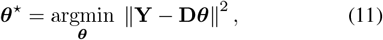

where observation matrices are

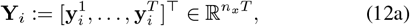

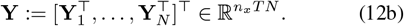

and design matrix

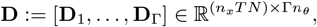

in which each 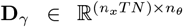 represents the block corresponding to type *γ* agents. The latter are defined as

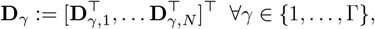

where *N* := | *L*| is the total number of cells in the dataset, considered here as a single time-lapse experiment.

For all the cells *I* ∈ *L*, we specify the entries of the observation matrices (12a) as

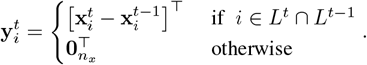

Instead, the single-cell design matrices are constructed as

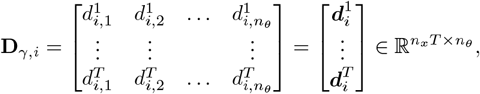

verifying that for each *t*

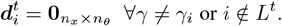

This condition, adding sparsity to the design matrix, ensures feasibility by limiting the data to cells present at the current frame *t* and by selecting different weights based on type *γ*_*i*_. Otherwise, the entries of cell *I ∈ L*^*t*^ are functions of the considered cell and its neighbor states

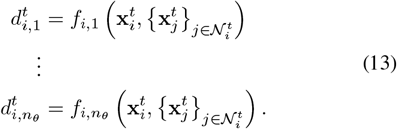

The solution of (11) that best fits cell trajectories extracted from a single multi-culture experiment is

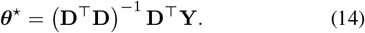

Note that while *n*_*x*_*TN* ≫Γ*n*_*θ*_ is necessary for the LS problem to be well-posed, invertibility of **D**^T^**D** additionally requires full column rank. This depends on the informativeness of the collected cell trajectories. For instance, if cells exhibit minimal motion, certain dynamical terms may remain insufficiently excited, potentially causing rank deficiency despite abundant data. To verify these conditions, we analyzed the design matrix constructed from our dataset. We confirmed that rank(**D**) = Γ*n*_*θ*_, ensuring full column rank. Specifically, we obtained a condition number of ≈ 4.76 10^3^, a value that is generally regarded as numerically safe for least-squares computations in double-precision arithmetic. To further improve numerical stability and reliability, we applied Cook’s distance metric [38] to identify and remove influential outliers, removing observations exceeding a threshold of 4*/* (*n*_*x*_*TN* − Γ*n*_*θ*_). The latter is a common heuristic for outlier detection in regression analysis. This outlier removal reduced the sum of squared residuals (SSR) from approximately 20.63 to 6.79, significantly enhancing both the fit quality and the robustness of the parameter estimates. Figure 2 illustrates the Cook’s distance values across samples, highlighting the detected outliers (3.6% of total data). Since at least one cell is present in every frame (*L*^*t*^ ≠∅ ∀*t*), all rows of the design matrix correspond to valid observations, thus ensuring consistency of the identification procedure.

**Figure 2.**
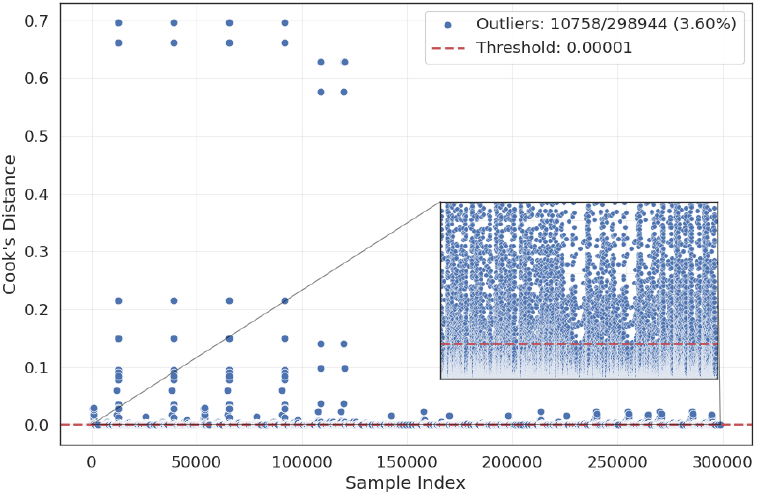
Cook’s distance values across samples, highlighting detected outliers (blue dots) based on a threshold (dashed red).

In the case of multi-experiment data, such as in our scenario, the process involves stacking observation vectors and design matrices for each data collection. Moreover, considering the scarcity of data in biological cell experiments, transformed versions of the same data sequences can be included to enhance robustness concerning invariance transformations, such as rotations, and horizontal and vertical flips. Details on the augmentation of the dataset are given in Section V on numerical experiments.

## Multi Cell Tracking Algorithm

In this section, we detail our solution to the cell-tracking problem outlined in Section II. Our Heterogeneous Open Multi-Cell EKF-based Tracking (HEOM-EKF) algorithm introduces a novel motion prediction step employing a tailored Extended Kalman Filter for Open Multi-Agent systems that leverages the models described in Section III. Moreover, our algorithm includes a procedure to estimate cell population changes and reconstruct the cell lineage tree. The main steps of the algorithm are detailed in Algorithm 1.

Here, the set estimations 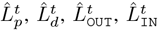 and 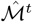 refer to the previous definitions of true quantities introduced in Section III defined in (1)-(3). We also provide a graphical illustration (see Figure 3) of how HEOM-EKF behaves in the presence of a mitosis event and how these label set estimates change accordingly across frames.

**Figure 3.**
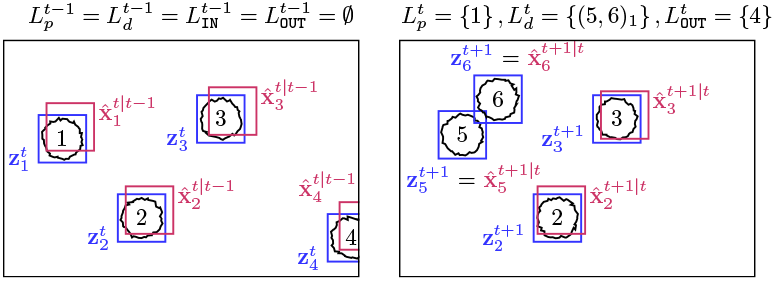
An illustrative example of label sets at different frames in a tracking scenario

**Figure 4.**
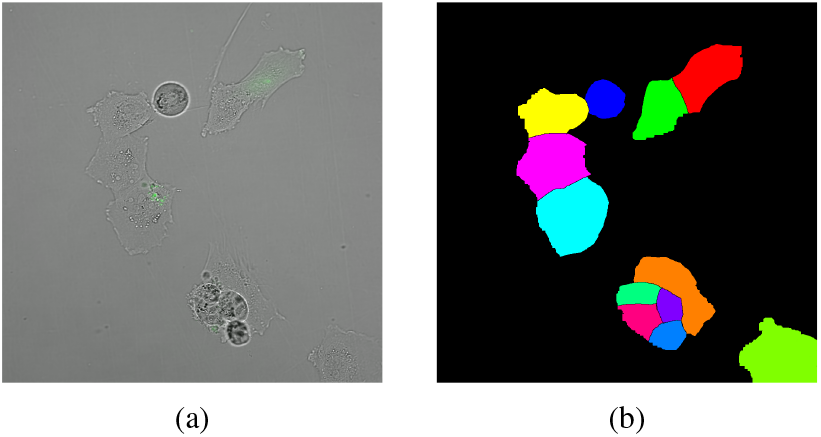
Comparison between the original microscope image (a) and its final segmented version (b).

In particular, the algorithm starts by initializing the state guess with detections at the first frame and setting up the EKF matrices. Then, throughout each frame, the algorithm iterates through three main steps. First, get the cell detections related to the current image. Second, it predicts the next state via the EKF prediction step leveraging the interaction dynamics learned from real data, whose identification of parameters is described in Section III-C. Third, it associates detections to predictions and updates the state estimate through the so-called EKF update step. Differently from standard filtering schemes, this step also takes into account changes in the number of cells. Specifically, when a cell population change is predicted at current frame, through MITOSIS-DETECT and IN-OUT-DETECT procedures (see Algorithms 2-3), the EKF update variables are adapted to account for dimensionality variations (see Algorithm 4). We now detail all the steps in the algorithm.

### Algorithm 1

HEOM-EKF

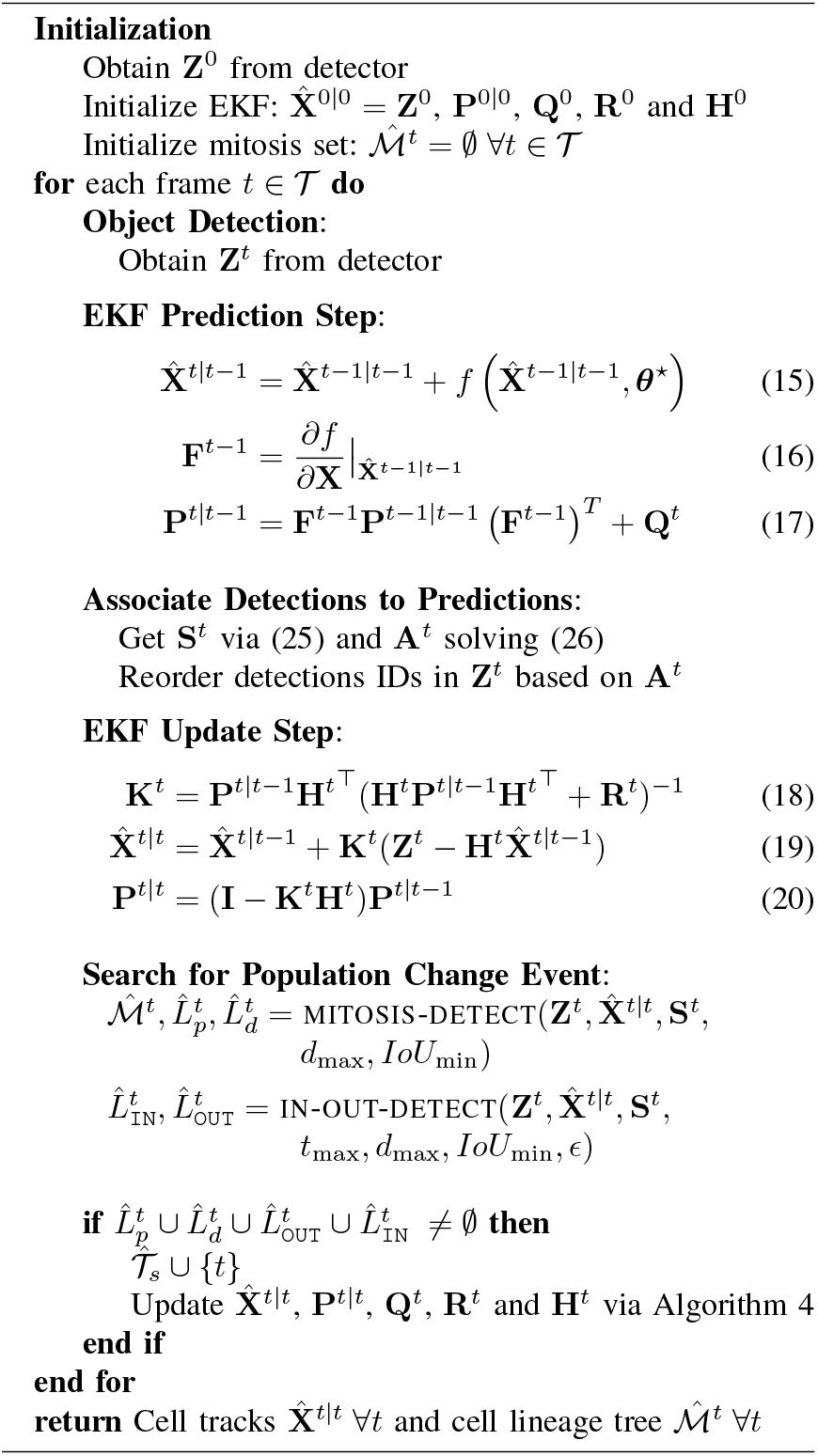

### A. Object Detection

The object detection phase serves as the initial step in our algorithm, with the primary goal of localizing cells in consecutive frames of the input video. All quantities and computations in our algorithm are directly based on the pixel coordinates of the detected cells. This ensures consistency with the spatial information provided during the object detection step. In particular, the measurement of cell *i* obtained via object detection at frame *t* is an oriented bounding box. Formally

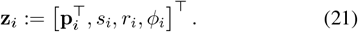

Here, **p**_*i*_ denotes the center coordinates of the bounding box for the *i*^th^ cell. The terms *s*_*i*_, *r*_*i*_ ∈ℝ represent the area and aspect ratio of the detected region, respectively, while *ϕ*_*i*_∈ [0, 2*π*] indicates the orientation angle.

It is worth noting that the object detection is performed independently on each frame, producing frame-specific cell detections and measurements, denoted as 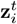 . The identity labels *i* are assigned arbitrarily by the detector, regardless of the method used. To ensure consistent tracking across frames prediction and data association steps (see Sections IV-C and IV-B) are needed to reorder and match cell identities over time.

### B. Motion Prediction

At each time *t*, the object detection phase yields a collection of measurement vectors 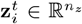 (cf. Section IV-A). Based on that, we can now detail the motion prediction step.

Let 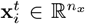 denote the true, unknown state of cell *i* at time *t*, represented in the format of equation (6). Accordingly, the detection process at each frame *t* can be modeled by the following measurement equation

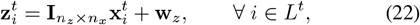

in which 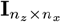 is the observation matrix, which is assumed to be constant and identical across all measurements. The term 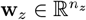 denotes zero-mean Gaussian noise with covariance **R**^*t*^.

Let us introduce 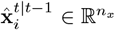 as the predicted state of *i*^th^ cell at time *t*, obtained after a prediction step in which the measurement **z**^*t*−1^ is known. Instead, 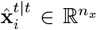 represents the updated estimate through the measurement 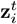 obtained after an estimation update step. To retrieve the aforementioned predictions, our tracking algorithm adopts an Extended Kalman Filter [39] incorporating as nonlinear dynamics the open multi-agent model introduced in Subsection III-B.

We first define

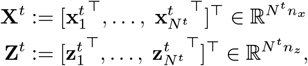

as the stack of cell population states and measurements. Similarly, we denote by 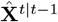 and 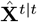 the stacks of predicted and updated state estimates, respectively. According to expression (22), the stack measurement vector is given by

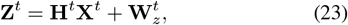

where 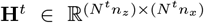 and 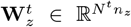 are the stacks ∀ *i* ∈ *L*^*t*^ of observation matrices and measurement noise vectors, respectively.

The multi-agent dynamics used to predict the *i*^th^ cell state belong to the class of models introduced in Section III-B, i.e.

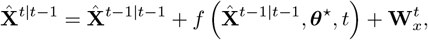

which is the predicted version of equation (5) written in compact form. In particular, 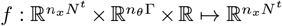 suitably stacks the dynamics in (8) for all the agents present at current time, and 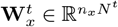 stacks the noise vector **w**_*x*_ for all the cells. The noise 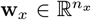 is a Gaussian random variable with zero mean and covariance **Q**. The optimal parameters in (15) are learned by solving (14) with properly constructed **D** and **Y** (see Section III-C) on real data. The step in (17) is the update of the so-called estimation error covariance matrix 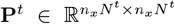 The steps in (18)– (20) instead perform the update steps of the EKF. Here, 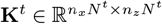 is the Kalman gain matrix.

Thanks to this motion prediction design, our tracker can operate on a unified state representation for the heterogeneous cell populations taking advantage of the sparse structure of the dynamics. Indeed, the motion prediction of each cell is performed considering interactions between neighboring cells rather than independent dynamical entities.

### C. Data Association

The data association step bridges the gap between detected cells across consecutive frames, preserving their identities over time. Indeed, given two consecutive time frames *t* − 1 and *t*, the *i*^th^ cell may be detected and associated with different labels. At the generic time *t*, the EKF predictions in (15) provide the estimate of a novel bounding box 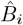 for cell *i*. This predicted box is centered in 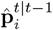 and has area and aspect ratio 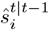 and 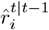, respectively. Considering the cell state as in (6), also the predicted box orientation 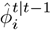 is present.

Similarly, let *B*_*j*_ denote the *j*^th^ bounding box found by the detection step. Let us introduce the similarity matrix 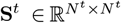 at image frame *t*, with single entry

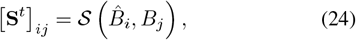

Where 𝒮 is a similarity function computed between the predicted bounding box of *i*^th^ cell and the detected one of *j*^th^ cell at time *t*. In this work, we use the so-called Intersection Over Union (IoU) metric [40], so that

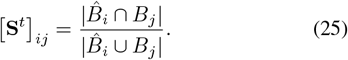

Given **S**^*t*^, the data association problem regards finding an optimal assignment matrix **A**^*t*^ at each frame of the video that maximizes the total similarity across all associations. This means solving at each time *t* the following linear-assignment problem

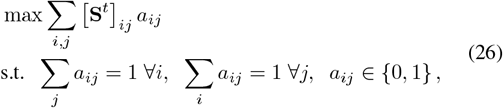

where each entry *a*_*ij*_ = 1 in **A**^*t*^ indicates that *j*^th^ detection is assigned to *i*^th^ prediction, and *a*_*ij*_ = 0 otherwise. This problem can be solved in polynomial time leveraging, e.g., the Hungarian algorithm [41]. To reject associations with a poor similarity index, we enforced a threshold for the IoU function in (25), denoted as *IoU*_min_.

We now define the set *𝒰* containing unmatched elements at time *t*, which can be further divided into unmatched predictions 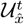 and unmatched detections 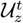 Specifically, the set of unmatches is defined as

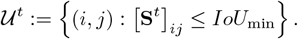

Then, we derive the subsets for unmatched predictions and unmatched detections

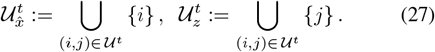

To provide better context, unmatched predictions and detections arise within a tracking algorithm where associations between predicted states and detections may not always be successful. In such cases, elements that fail to match are categorized into these sets to facilitate further analysis or handling in the tracking process.

### D. Cell Population Changes and Cell Lineage Tree

In this subsection, we define three routines used in our cell tracking strategy HEOM-EKF. We first introduce a method for estimating cell mitosis events (Algorithm 2). Then, we describe how to detect the entry and exit of cells within the microscope FOV (Algorithm 3). Lastly, we present the procedure to update the dimensions of the EKF matrices in response to the changes in the cell population (Algorithm 4).

We begin by introducing some quantities used within Algorithm 2 and 3. Let 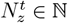 be the number of detected cells and 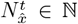 the number of predicted cell states. Also, let 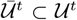 be the subset of unmatched predicted and detected cells whose bounding boxes have no intersection and whose distance exceeds the mitosis threshold *d*_max_. This distance is determined based on real observed behaviors within the coculture experiments, and it represents the maximum center distance at which a mitosis event is usually detected. Formally,

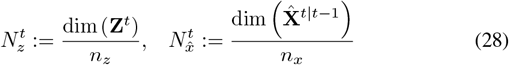

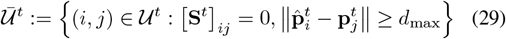

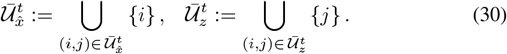

The ability to accurately determine mitosis events is important for extracting relevant information about the evolution of heterogeneous cell populations over time. A possible approach consists of constructing the cell lineage tree for each cell. To this end, in the following, we present Algorithm 2.

The proposed method estimates mitosis occurrences at a given frame *t* by computing the set estimates 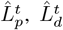, and thus 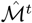 . In particular, the algorithm leverages the sets in (27) and (30), together with the quantities in (28), to distinguish new detections due to mitosis, cells entering the camera FOV or unmatched detections. After tracking is completed across the entire video sequence, the lineage of each cell is constructed using the parent-child relationships stored in the sets 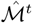 for all *t*∈ 𝒯 . Note that 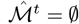 at frames where no mitosis events are detected.

We now present Algorithm 3 to estimate the occurrence of cells entering or exiting from the camera FOV.

#### Algorithm 2

MITOSIS-DETECT : Get set estimate 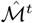

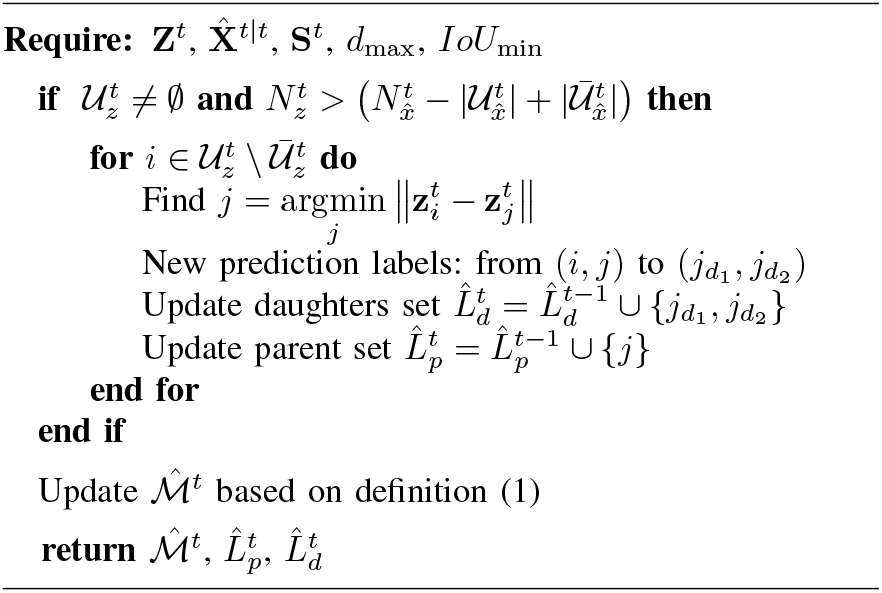

#### Algorithm 3

IN-OUT-DETECT : Get set estimates 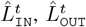

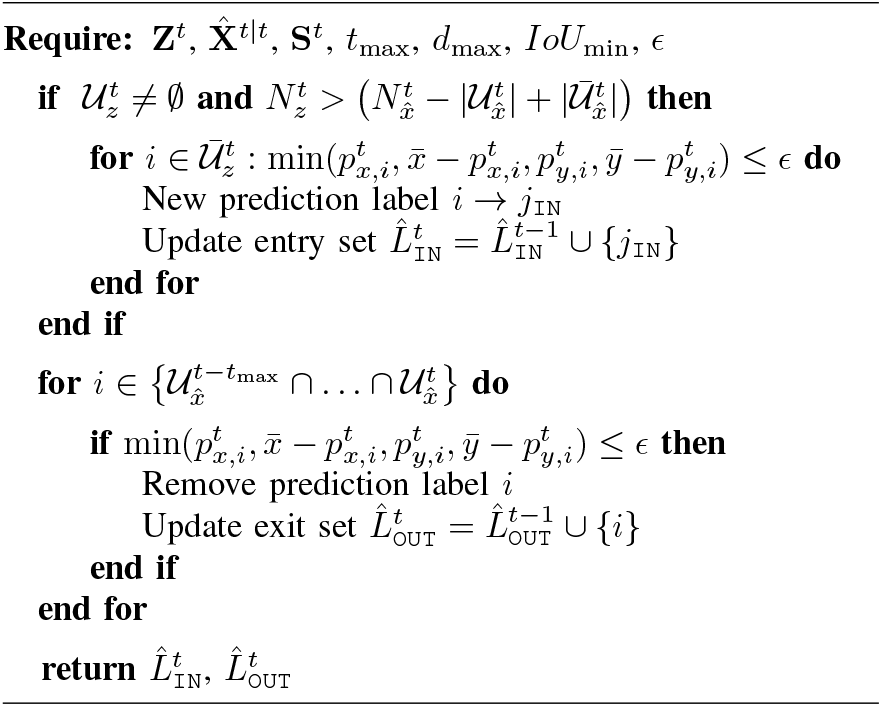

Analogously to Algorithm 2, this method begins by checking for unmatched detections at the current frame *t*, using the sets defined in (27) and (30), along with the values in (28). It then examines the set 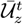 to identify detected cells that are *ϵ*-close to the image boundaries 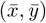, i.e., cells that have recently entered the field of view. To detect cells that are leaving the microscope FOV, the algorithm instead looks for predictions that remained unmatched over the past *t*_max_ time steps. If these predictions are also *ϵ*-close to the image boundaries, they are considered as outcoming cells.

Algorithm 4 takes care of adapting the dimensions of EKF matrices. Specifically, it reorganizes the terms 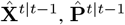, **H**^*t*^, **R**^*t*^, and **Q**^*t*^ to ensure feasible tracking. Especially, let us define the functions used for including new daughter cells into the tracking procedure as

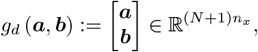

in which 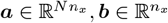 Also, let

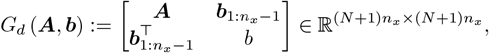

in which 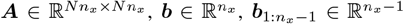 and *b ∈ ℝ*. Instead, the functions used to remove a parent cell within HEOM-EKF are

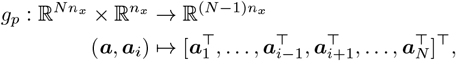

in which ***a***_*i*_ belongs to 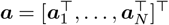, and

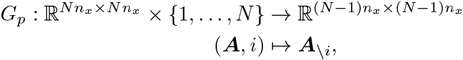

with ***A***_*\i*_ being the matrix ***A*** with the *i*^th^ row and column removed.

#### Algorithm 4

Update EKF dimensionality

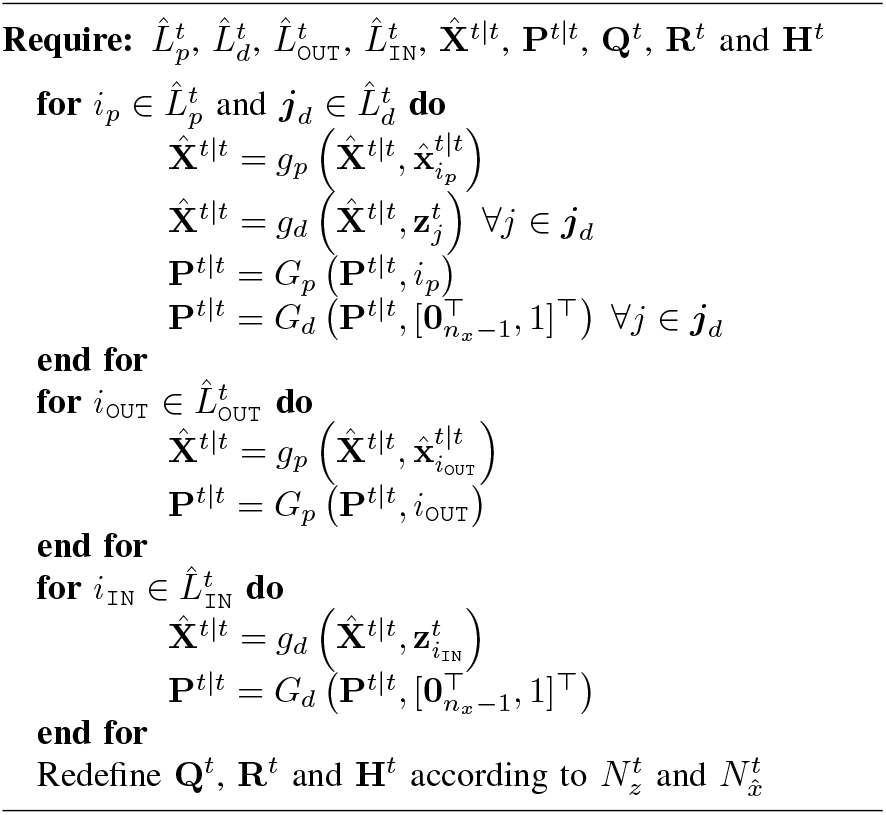

## V. Numerical Experiments

In this section, we first introduce the dataset used in our simulations. Then, we present the results of the proposed cell-tracking algorithm, analyze the optimal multi-agent model parameters that confirm cell type discrimination, and discuss the extracted cell lineage tree.

### A. Dataset

The dataset used in this study was derived from raw time-lapse image sequences captured using a confocal microscope (A1R MP microscope, Nikon) at IRCCS Istituto Ortopedico Rizzoli. The human 143b line were obtained from the American Type Cell Culture Collection (ATCC) and were cultured in Minimum Essential Medium Eagle Alpha modification (Sigma) with 10% fetal bovine serum (FBS, Sigma) and 1% penicillin/streptomycin (P/S) (Euroclone) and used for co-culture experiments after trypsinization and semi-confluence. Human adipose-derived MSC (ATCC) were maintained in complete alpha-MEM with 10% fetal bovine serum (FBS, Sigma) and 1% P/S and used for co-culture experiments after trypsinization and semi-confluence. Cells were incubated at 37 °C in 5% CO_2_. GFP-transfected MSC cells, obtained as previously described [31], were used in co-culture experiments. For co-cultures, 143b cells and GFP+ MSC were mixed 1:3 and seeded at low density in complete alpha-MEM in petri dishes for time-lapse imaging. The transfection with GFP allowed us to clearly distinguish MSC from 143b cells and, consequently, accurately assign the measured features between the two cell types. The set of experiments carried out consists of 4 videos with 172 frames each with images captured every 4 minutes (25x immersion objective, NA 1.1, RI 1.333). Initial processing of the raw images was performed by using NIS Elements AR 5.40.01 software (Nikon), employed on all the images to perform semantic segmentation, labeling, and classification by cell type. We performed these tasks by combining manual adjustments with automated image processing tools. In particular, we first manually segmented representative cell regions and then used the software to automatically propagate and refine the contours across subsequent frames by tuning intensity thresholds and edge-detection parameters. When cells overlapped or formed dense clusters, segmentation was instead carried out entirely by manual annotation. Since the process could be subject to human and/or software automation errors, we refer to these annotations as the gold standard, rather than ground truths. Following segmentation, the software extracted around 70 features per segmented object, including morphological and positional properties, allowing for detailed temporal analysis of each cell. These features, after filtering only the ones relevant to our purposes, provided a comprehensive characterization of the cells’ dynamics and their interactions over time. To refine the dataset further, additional image processing techniques were applied, focusing on denoising and fixing any labeling or segmentation errors. The latter postprocessing corrections ensured enhanced data quality. Before presenting the final results, we report summary statistics of key cell features extracted from the dataset (Figure 5). These metrics reveal clear morphological differences between MSC and 143b cells and motivate the design of the proposed open multi-agent dynamics. Although perimeter, circularity, and elongation are not explicitly modeled as state variables, their morphological information is implicitly encoded in the bounding box representation. In particular, the aspect ratio *r*_*i*_ provides a proxy for circularity and elongation, capturing type-specific shape characteristics. The target aspect ratio 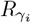 for each cell type is derived from the mean circularity observed in the training data, thereby embedding morphological priors into the tracking framework. As expected, MSC display a more elongated, fibroblast-like morphology, whereas osteosarcoma (143b) cells tend to be more rounded. On average, MSC are larger than 143b cells in terms of both area and perimeter.

**Figure 5.**
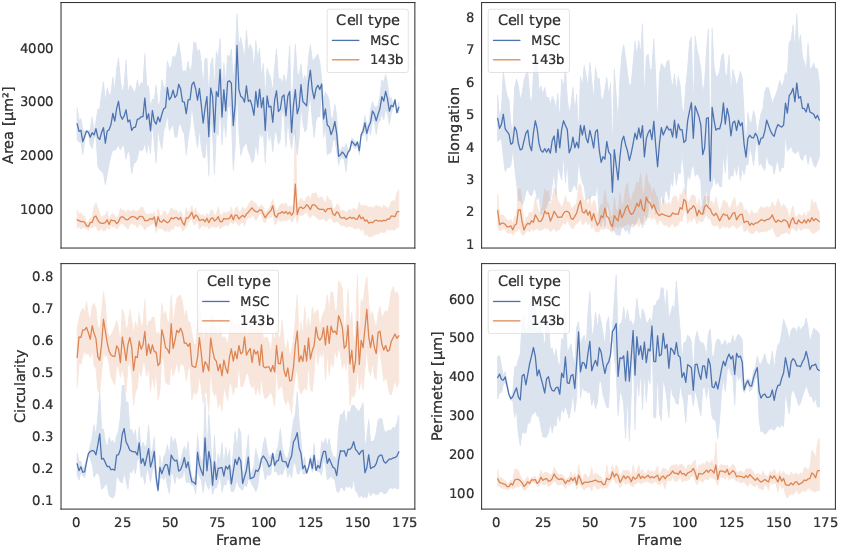
Mean and variance over time for the area, outer perimeter, circularity, and elongation, categorized by cell type.

Finally, we want to highlight that this dataset is a unique resource to study cellular behavior in co-culture frameworks, setting it apart from most publicly available datasets, such as the ones from the Cell Tracking Challenge, which only focus on monoculture experiments.

### B. Cell Detection in Video Frames

In our experiments, we adopt a semi-automatic approach using the proprietary image-analysis software included with our microscope. This choice offers several advantages: (i) it simplifies the detection pipeline, allowing us to focus on assessing the performance of our novel motion prediction module; (ii) it avoids the need for large, annotated datasets typically required for supervised training; and (iii) it ensures compatibility and consistency with the data acquisition process, as the detection tool is tailored to the imaging system. For each video frame, the software provides region proposals (cell segmentations), from which axis-aligned or oriented bounding boxes can be extracted using basic image processing techniques. State-of-the-art cell segmentation tools rely on training and deploying a dedicated deep learning model for cell detection, as in [42], [43]. However, these approaches typically require a large amount of data and do not discriminate between different cell populations. Our multi-agent approach to cell tracking aims to introduce dynamical system knowledge into the process. This improves tracking performance even when the object detection component cannot exploit a large amount of data.

### C. Experimental Results

We implemented the proposed methodology via the Python 3 programming language, testing our algorithm on co-culture videos from *in vitro* experiments involving MSC and osteosarcoma cells. Given the limited dataset of only four videos, we employed a cross-validation approach [44]. Specifically, we trained the models described in Section III, by computing the optimal solution ***θ***^∗^ as in (11) considering three videos regarding a co-culture scenario (Γ = 2). Then we evaluated the tracking performances on the remaining one. We repeated this procedure for all possible combinations of training and test videos. To improve the robustness of our estimation and mitigate overfitting due to the small dataset, we adopted data augmentation techniques [45]. Specifically, we applied rotations of angles *α* ∈90^°^, 180^°^, 270^°^ and orthogonal flips to each segmented image to then extract the corresponding rotated versions of 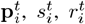 and 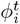 for all cells *i* at each frame *t*. After computing the required time derivatives, these transformed values were used to enlarge **Y** and **D** on the LS problem. Moreover, we worked on normalized data sequences to ensure better-conditioned matrix inversion, prevent feature predominance, and enhance the interpretability of parameters. The data augmentation and the subsequent identification process were performed offline. The model constants in equations (10), specifically the target area 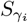 and aspect ratio 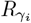 with *γ*_*i*_ ∈{MSC, 143b}, were derived directly from dataset statistics regarding mean area and elongation observed across cell trajectories over time (refer to Figure 5). We further analyzed the learned model parameters by examining their distributions across different training sets. This allowed us to evaluate if our open multi-agent model could distinguish between the two cell types (MSC and 143b cells) by capturing their distinct behaviors. The ability to automatically distinguish cells in a co-culture model, such as MSC and osteosarcoma cells, is crucial, as both originate from the same mesenchymal progenitor line and appear morphologically similar under direct microscope observation. Consequently, identifying and analyzing cellular behaviors and interactions between these two cell types to gain new insights into cancer progression and aggressiveness is currently limited to systems that are expensive, complex, and often impractical to use. Specifically, we examined the parameter sets obtained from training on all possible combinations of the four videos, i.e., the power set of the dataset, which consists of 2^4^ different training sets. For each training set, we computed the identified parameters and analyzed their statistical distribution using kernel density estimation (KDE) to obtain a smooth approximation of the underlying data distribution. Figure 6 illustrates the KDE-based distribution of the identified parameters for the two cell types across different training sets. The analysis reveals distinct parameter distributions, supporting the hypothesis that the proposed model effectively differentiates MSC from 143b cells.

**Figure 6.**
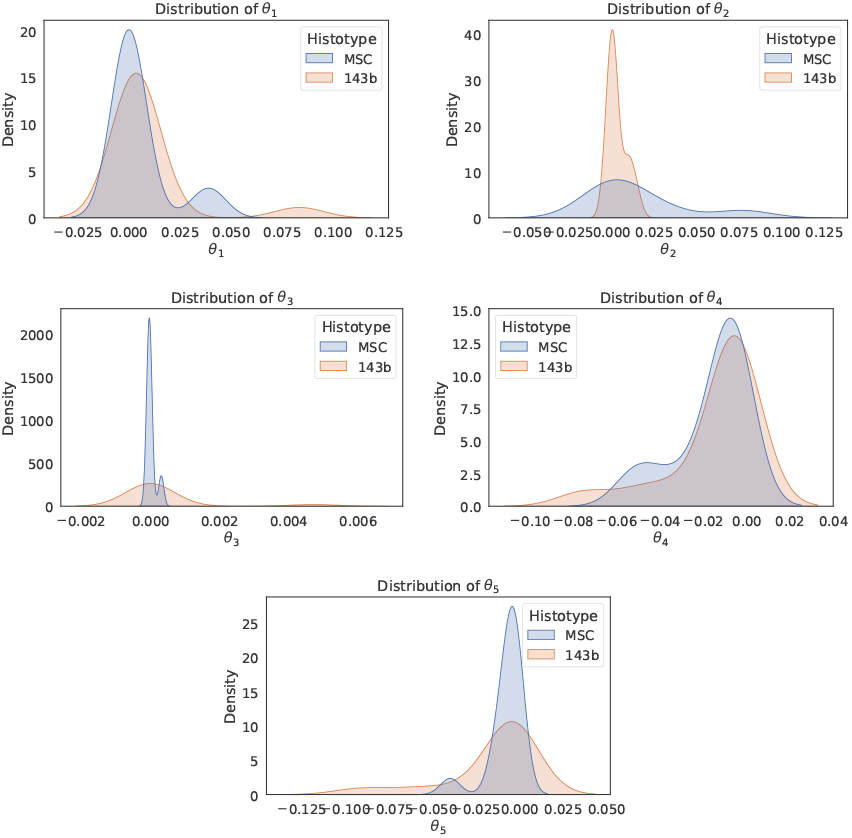
Distributions of the identified model parameters for MSC and 143b cells across diverse training sets.

From a biological perspective, the identified parameter weights can be interpreted as follows. The cohesion parameter 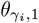 quantifies the tendency of cells to remain in proximity to their neighbors. It reflects the propensity of cells to lose cohesion, detach from the tissue, and migrate to distant sites, a phenotype commonly associated with tumor cells and increased aggressiveness and metastatic potential. In line with this interpretation, our results indicate that mesenchymal stromal cells (MSCs) exhibit a lower tendency to lose cohesion than tumor cells. The alignment parameter 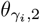 measures the tendency of cells to migrate in the same direction as their neighbors. The lower values with smaller variance observed in tumor cells suggest a more heterogeneous and less coordinated migratory behavior compared with MSCs, consistent with the disorganized migration typical of cancer cells that have lost responsiveness to local cues required for coordinated tissue dynamics and structural integrity. Further support for these findings is provided by the repulsion parameter 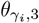 . Tumor cells display a greater tendency to distance themselves from neighboring cells than normal cells, indicating a disruption of the interaction mechanisms characteristic of organized tissues. Finally, the morphological parameters 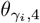 and 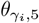, which regulate the tendency to maintain the cell-type-specific target area 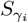 and aspect ratio 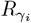, show similar distributions between the two cell types. MSCs maintain elongated and relatively stable morphologies around their characteristic shape, whereas 143b cells exhibit increased variability around their own, more rounded morphology, consistent with their enhanced motility and well-documented morphological plasticity. Overall, these results demonstrate that the identified parameters not only distinguish MSCs from 143b cells statistically, but also capture impaired regulation of cell-cell interactions. As such, they represent informative indicators of phenotypic differences between normal and tumor cells and may serve as indirect measures of tumor aggressiveness in biological assays, even in the presence of normal cells capable of modulating tumor behavior.

To further detail the tracking algorithm, we now describe the initialization and tuning of the EKF terms used within Algorithm 1. In our implementation of the Extended Kalman Filter, the error covariance matrix **P**^0|0^ is initialized by assigning higher variance values to state components that are initially unobserved, reflecting our uncertainty in these estimates. Instead, the measurement covariance matrix **R** and the process covariance matrix **Q** are set based on baseline estimates obtained from preliminary noise analyses of the dataset. To account for different noise conditions, we further explored scaled versions of these baselines using factors of 0.5, 1.0, and 2.0. This systematic exploration enabled us to quantify the impact of these initialization choices on tracking performance and so identify the optimal settings for our application.

We want to underline that, since our focus is on evaluating the performance metrics of our tracker rather than the detector, we conducted experiments by simulating the detection process using gold standard annotations with injected Gaussian disturbances. Specifically, the measurement errors were simulated considering random noise with zero mean and defined covariance for each annotation. In the following, we present some results considering the state format (7) since, being the standard detection format, it enabled us to perform a comparison with other methodologies. Figure 7 shows a video snapshot regarding the numerical experiments on one of the co-culture videos. Here, we display both the bounding boxes predicted by our algorithm and the original gold standard annotations.

**Figure 7.**
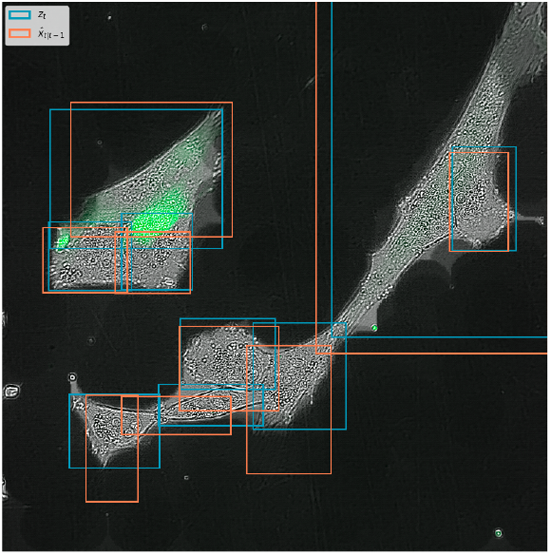
Predicted bounding boxes (orange) and gold standard annotations (blue) in a co-culture video frame.

We compared our scheme against two baseline multi-object tracking methods: SORT [46] and DeepSORT [47]. Both use a Kalman filter for motion prediction and the Hungarian algorithm for data association. SORT performs association using IoU only, whereas DeepSORT employs a neural network trained for re-identification to extract appearance features to improve association. For a fair comparison, the DeepSORT appearance model was fine-tuned on the images from our co-culture dataset. We initialized SORT and DeepSORT with the same matrices as in Algorithm 1 and simulated the detection process for our proposed solution. The chosen metrics are the Multiple Object Tracking Accuracy (MOTA), Multiple Object Tracking Precision (MOTP) [48] and Identity F1 Score (IDF1). We also report the average per-frame processing time, denoted as *T*_avg_ ∈ ℝ_*>*0_ for each algorithm. MOTA depends on the number of misses, false positives, and mismatches. MOTP compares the predicted bounding boxes against the real ones. IDF1 measures the accuracy of identity assignment by comparing correctly matched detections to the average number of ground-truth and predicted detections. Higher values of these metrics indicate better tracking performance. Table I summarizes the results of performing the cross-validation procedure on different experiments, referred to as “Well”. This comparison highlights the effectiveness of our approach, as evidenced by a good MOTA score, which reflects the algorithm ability to track cells accurately over time. Furthermore, the MOTP metric demonstrates that the proposed tracking model outperforms SORT and DeepSORT, providing more precise predictions. Note that, DeepSORT does not improve over SORT. We attribute this to scarcity of data: the appearance network in DeepSORT is a relatively large model, whereas our total dataset consists of only four videos. Fine-tuning such a network on so few sequences is unlikely to generalize and can even degrade association performance compared to a purely geometry-based method. Moreover, Table I reports the results of an ablation study that isolates the impact of the motion-prediction component. Indeed, HEOM-EKF, SORT, and DeepSORT share the same Track-by-Detection structure and a KF-based tracking scheme; the key difference lies in the open system model and relative enhanced version of EKF. In HEOM-EKF, this model is identified from real co-culture data and explicitly tailored to heterogeneous, time-varying cell populations, whereas SORT and DeepSORT rely on standard single-object kinematic models. Keeping detection and association fixed isolates the contribution of the proposed multi-agent motion model within the EKF prediction step. In terms of per-frame computation time, our method is slower than SORT but substantially faster than DeepSORT. This outcome is expected since, relative to SORT, HEOM-EKF employs a more sophisticated motion model and an EKF whose state dimension scales with the number of cells (while avoiding the computationally intensive appearance feature extraction required by DeepSORT). Despite this increased complexity, the per-frame runtime remains on the order of tens of milliseconds, which is negligible relative to the minute-scale image acquisition intervals typical of cell culture experiments, thereby ensuring compatibility with real-time microscopy.

**Table I.** Performance comparison.

| Metric | Experiment | SORT | DeepSORT | Alg. 1 |
| --- | --- | --- | --- | --- |
| MOTA | Well 1 | 74.23 % | 55.30 % | **83.56 %** |
|  | Well 2 | 40.78 % | 52.77 % | **67.58 %** |
|  | Well 3 | 76.38 % | 54.09 % | **87.02 %** |
|  | Well 4 | 79.97 % | 55.63 % | **96.17 %** |
| MOTP | Well 1 | 93.68 % | 85.33 % | **96.72 %** |
|  | Well 2 | 91.45 % | 84.04 % | **95.51 %** |
|  | Well 3 | 93.34 % | 83.56 % | **96.03 %** |
|  | Well 4 | 94.33 % | 85.38 % | **97.18 %** |
| IDF1 | Well 1 | 62.61 % | 50.31 % | **69.32 %** |
|  | Well 2 | 54.20 % | 38.79 % | **66.67 %** |
|  | Well 3 | 65.59 % | 47.76 % | **71.07 %** |
|  | Well 4 | **82.02 %** | 48.12 % | **80.88 %** |
| $T_{\text{avg}}$<br>[ms/frame] | Well 1 | **2.33** | 204.98 | 59.73 |
|  | Well 2 | **0.94** | 94.07 | 5.33 |
|  | Well 3 | **3.32** | 184.42 | 60.72 |
|  | Well 4 | **1.76** | 170.57 | 40.27 |

Additionally, the proposed algorithm predicts changes in cell population over time and, at the end of the time-lapse sequence, returns the estimated cell lineage tree 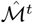 for the frames *t*∈{ 1, …, 172} of the current *in vitro* experiment. Figure 8 shows a prediction result of the latter. Here, the state format in (6) was preferred since it allowed for more precise type discrimination and parent-daughter relation prediction. The depicted example of an estimated cell lineage tree is a reliable result despite the absence of a true reference family tree for direct comparison. In fact, MSC cells rarely divide (considering the duration of the single experiment of about 10 hours), while 143b osteosarcoma cells, due to their oncogenic nature, exhibit a significantly higher mitosis rate. The predicted lineage tree captures these expected proliferation patterns, reinforcing its plausibility.

**Figure 8.**
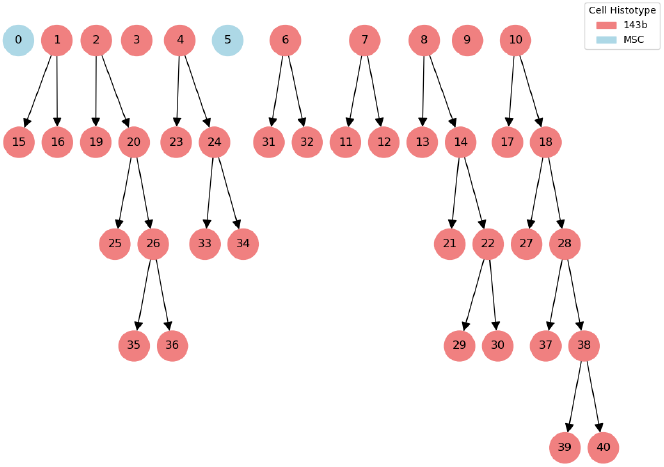
Example of a predicted cell lineage tree.

The proposed framework shows promising performance, but the limited dataset size (4 co-culture videos) must be acknowledged. Despite cross-validation and data augmentation, overfitting risks remain. This limitation reflects a broader challenge in cancer-biology: high-quality time-lapse datasets of co-cultures are scarce due to experimental complexity and cost. Thus, the proposed work represents a proof-of-concept in this data-scarce context and highlights that our proposed method can represent a valuable solution as opposed to Machine Learning approaches that typically require large datasets for training. Moreover, this study provides a unique dataset for investigating MSC-143b interactions and evaluating tracking in co-culture experiments.

## VI. Conclusions

In this paper we developed a novel algorithm for tracking cells in optical microscopy videos through open multi-agent modeling. System identification techniques applied to *in vitro* experiments enabled to learn cell type-specific behaviors, accounting for cell-to-cell interactions even in multi-culture scenarios. The open multi-agent system framework is the building block of the proposed cell tracking algorithm that is based on a generalized EKF. Cross-validation tests with proper performance metrics were shown together with a comparison with two baseline algorithms in multi-object tracking. Results demonstrate that the proposed method represents a valuable solution in a data-scarce context as cancer biology for tracking heterogeneous cell types, capturing cell-to-cell interactions and generating the cell lineage tree. This study establishes a unique testing framework using a dataset obtained from *in vitro* co-culture experiments with tumor and normal cells.

---

**Andrea Tramaloni** (S’23) received a master’s degree in automation engineering from the University of Bologna in 2022. Currently, he is a Ph.D. student at the Center for Research on Complex Automated Systems (CASY), Department of Electrical and Information Engineering (DEI), University of Bologna, Bologna, Italy. He was a visiting scholar at the University of Bristol in 2025. His research interests include systems biology, cybergenetics, multiagent systems, and optimal control.

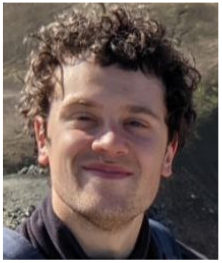

**Andrea Testa** received the Laurea degree “summa cum laude” in Computer Engineering from the Università del Salento, Lecce, Italy in 2016 and the Ph.D degree in Engineering of Complex Systems from the same university in 2020. While working on this paper, he was a Junior Assistant Professor at Alma Mater Studiorum Università di Bologna, Bologna, Italy. He was a visiting scholar at LAAS-CNRS, Toulouse, (July to September 2015 and February 2016) and at Alma Mater Studiorum Università di Bologna (October 2018 to June 2019). His research interests include control of UAVs and distributed optimization. Dr. Andrea Testa served as an Associate Editor for the *19th* and *20th IEEE International Conference on Automation Science and Engineering (CASE)*.

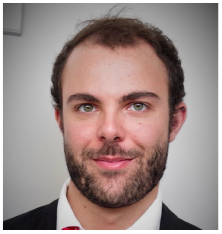

**Sofia Avnet** received a Degree in Medical Biotechnologist, in 2000, and the Ph.D. degree, in 2005, a second Degree in cellular and molecular biology, in 2010, and the Specialization degree in pharmacology and toxicology, in 2023. During the first master’s degree, she began research on bone tumors with the Kimmel Cancer Institute and the IRCCS Istituto Ortopedico Rizzoli. For the Ph.D. degree, she continued studying biological mechanisms of bone cancers with the Institute for Cancer Research, Candiolo. She also conducted research on bone pathophysiology as a Visiting Researcher with McGill University, in 2006, and a Research Fellow with the University of Turku, in 2007. From 2008 to 2019, she was a Biotechnologist with the Rizzoli Orthopedic Institute, focusing her research on tumor invasiveness, chemoresistance, microenvironment, and metabolism in bone cancers. Since 2019, she has been an Assistant Professor with the University of Bologna, where she develops 3D bone tissue and bone cancer models using spheroids, microfluidics, and bioprinting. Over her career, she has received several research grants as a Principal Investigator from the Italian Association of Cancer Research and the Ministry of Health. Dr. Avnet has been the President of the International Society of Cancer Metabolism (2015-2016).

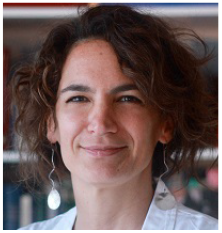

**Stefania Massari** holds an MSc in Medical Biotechnology (2022) and a BSc in Biological Science and Technology (2018), both from the University of Trieste. She has been a biotechnologist and research fellow at the Rizzoli Orthopaedic Institute, where she has worked on molecular oncology and cancer metabolism. She has contributed to an AIRC-funded project investigating the effects of tumor microenvironment acidity on lipid metabolism in osteosarcoma. Currently, she is a PhD student at the Medical University of Vienna.

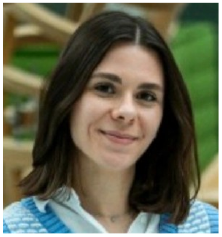

**Gemma Di Pompo** is a biologist (Master Degree in 2009 from University of Bologna), with a PhD in Biotechnology, Pharmacology, and Toxicology (2013) from the University of Bologna. In 2014, she has been a visiting researcher at Orofacial Development and Regeneration Unit (Institute of Oral Biology, Centre of Dental Medicine, University of Zurich, Switzerland). Currently, a senior researcher at the Rizzoli Orthopaedic Institute, her work focuses on bone biology, cancer microenvironment, and regenerative medicine. She has participated in multiple national and international research projects on bone tumors, cancer-associated pain, and advanced therapies using 3D bioprinting and tissue engineering. She was recently awarded a research grant as Principal Investigator by the AIRC Foundation to support her ongoing studies in targeting nerve-cancer crosstalk as a novel approach to improve osteosarcoma treatment. She has authored 31 peer-reviewed publications and has received recognitions, including the European Orthopaedic Research Society (EORS) Exchange Travel Grant and the “Top Junior Scientist” award.

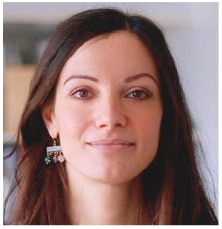

**Nicola Baldini** received the Degree in medicine and surgery, the Certification degree in orthopaedics, and the Certification degree in oncology from the University of Bologna, Bologna, Italy, in 1981, 1987, and 1990, respectively. From 1988 to 1989, he was a Research Fellow with Harvard University. From 2013 to 2018, he was a Visiting Professor with Kyoto Prefectural University of Medicine. He is currently a Professor with the Department of Biomedical and Neuromotor Sciences, Alma Mater Studiorum, University of Bologna, and the Director of the Department of Research, Innovation and Technology, IRCCS Istituto Ortopedico Rizzoli, Bologna. His research interests include the microenvironment of bone tissue under physiological and pathological conditions, bone cancers, and regenerative medicine. Prof. Baldini has served as the President for the Connective Tissue Oncology Society, from 1999 to 2000, the European Orthopaedic Research Society, from 2008 to 2012, and the International Society for Cancer Metabolism, from 2019 to 2020.

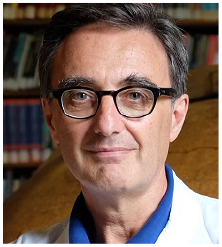

**Giuseppe Notarstefano** (M’11) is a Professor in the Department of Electrical, Electronic, and Information Engineering G. Marconi at Alma Mater Studiorum Università di Bologna. He was Associate Professor (June ‘16 - June ‘18) and previously Assistant Professor, Ricercatore, (February ‘07 - June ‘16) at the Università del Salento, Lecce, Italy. He received the Laurea degree “summa cum laude” in Electronics Engineering from the Università di Pisa in 2003 and the Ph.D. degree in Automation and Operation Research from the Università di Padova in 2007. He has been visiting scholar at the University of Stuttgart, University of California Santa Barbara and University of Colorado Boulder. His research interests include distributed optimization, cooperative control in complex networks, applied nonlinear optimal control, trajectory optimization and maneuvering of aerial and car vehicles. He serves as an Associate Editor for IEEE Transactions on Automatic control, IEEE Transactions on Control Systems Technology and IEEE Control Systems Letters. He is also part of the Conference Editorial Board of IEEE Control Systems Society and EUCA. He is recipient of an ERC Starting Grant.

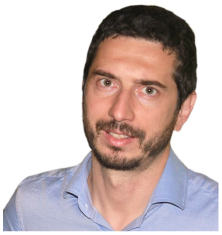

